# ABHD6 selectively controls metabotropic-dependent increases in 2-AG production

**DOI:** 10.1101/2022.05.18.492553

**Authors:** Simar Singh, Dennis Sarroza, Anthony English, Dale Whittington, Ao Dong, Mario van der Stelt, Yulong Li, Larry Zweifel, Michael R. Bruchas, Benjamin B. Land, Nephi Stella

**Affiliations:** Departments of Pharmacology, University of Washington, Seattle, USA; Medicinal Chemistry, University of Washington, Seattle, USA; Peking University School of Life Sciences, PKU-IDG/McGovern Institute for Brain Research, Peking-Tsinghua Center for Life Sciences, Academy for Advanced Interdisciplinary Studies, Peking University, Beijing, China; Leiden Institute of Chemistry, Leiden University, Leiden, Netherlands; Departments of Psychiatry and Behavioral Sciences, University of Washington, Seattle, USA; Departments of Center for the Neurobiology of Addiction, Pain, and Emotion, University of Washington, Seattle, USA; Departments of Center for Cannabis Research, University of Washington, Seattle, USA; Departments of Anesthesiology and Pain Medicine, University of Washington, Seattle, USA

## Abstract

The most abundant endocannabinoid (**eCB**) in the brain, 2-arachidonoyl glycerol (**2-AG)**, is hydrolyzed by α/β-hydrolase domain containing 6 (**ABHD6**); yet how ABHD6 controls stimuli-dependent increases in 2-AG production is unknown. To explore this question, we leveraged the recently developed 2-AG sensor, GRAB_eCB2.0_, and found that stimulation of Neuro2a cells in culture with bradykinin (**BK**) acting at metabotropic B_2_K receptors and ATP acting at ionotropic P2X_7_ receptors led to differential increases in 2-AG levels. B_2_K triggered increases in 2-AG levels via diacylglycerol lipase (**DAGL**), and this mechanism was potentiated by increases in intracellular calcium and ABHD6 inhibition. By contrast, P2X_7_-triggered increases in 2-AG levels were dependent on DAGL and extracellular calcium but unaffected by ABHD6 inhibition. Thus, ABHD6 preferentially regulates metabotropic-dependent increases in 2-AG levels over ionotropic-dependent increases in 2-AG levels. Our study indicates that ABHD6 selectively controls stimuli-dependent increases in 2-AG production and emphasizes its specific role in eCB signaling.

## Introduction

eCB signaling plays a fundamental role in multiple physiological processes throughout the body, including in cannabinoid receptor type 1 (**CB**_**1**_**R**)-dependent regulation of neurotransmitter release, neuronal metabolism, and neuronal phenotypes^1,2^. The two best-studied eCBs, arachidonoylethanolamide (anandamide, **AEA**) and 2-AG, differ in at least 4 key aspects: 1] their biosynthetic pathways (NAPE-PLD and DAGL, respectively), 2] their relative abundance in cells (AEA often 10-1000 fold less abundant than 2-AG), 3] their potency and efficacy at CB_1_R (AEA acts with high potency and as partial efficacy agonist, whereas 2-AG acts with lower potency and as full efficacy agonist), and 4] their enzymatic inactivation (FAAH hydrolyzing AEA, and MAGL and ABHD6 hydrolyzing 2-AG). How MAGL controls stimuli-dependent increase in 2-AG production and activity at CB_1_R has been extensively studied^3^; yet how ABHD6 controls stimuli-dependent increase in 2-AG production and activity at CB_1_R remains poorly understood^4^. Specifically, both enzymes hydrolyze 2-AG to form glycerol and arachidonic acid, and measures of their enzymatic activity in brain homogenates show that MAGL accounts for approximately 80% of total 2-AG hydrolyzing activity and ABHD6 only for 5-10% of total 2-AG hydrolyzing activity^5^. However, evidence gathered with intact brain tissues and cells in culture show that the 2-AG hydrolysis measured in homogenates cannot account for the relative involvement of each enzyme in the control of 2-AG’s activity at CB_1_R; rather, their subcellular expression and their proximity to 2-AG production and CB_1_R in intact tissue better reflects their relative contribution to the control of 2-AG-CB_1_R signaling^4^. For example, ABHD6 inhibition does not influence 2-AG tone in mouse cortical slices and yet it enables the establishment of LTD triggered by subthreshold stimulation to the same extent as MAGL inhibition^6^. Furthermore, ABHD6 inhibition regulates short term plasticity in cell culture model systems to the same extent as MAGL inhibition^7^. These studies indicate that ABHD6 mainly controls 2-AG-CB_1_R signaling in neurotransmitter-stimulated neurons; however, whether ABHD6 inhibition similarly influences distinct stimuli-dependent increases in 2-AG production and subsequent activity at CB_1_R is unknown. Understanding how ABHD6 controls eCB signaling is critical towards gaining insights into its role in eCB signaling and to help develop the therapeutic properties of ABHD6 inhibitors.

Genetically encoded fluorescent sensors were recently engineered and allow for real-time detection of changes in the levels of select neurotransmitters and neuromodulators^8^. This technology leverages the high binding of endogenous ligands to specific receptors, which stabilizes a conformation to elicit fluorescence of a circularly permutated-green fluorescent protein (**cpGFP**) introduced in the third intracellular loop of the G protein-coupled receptors (**GPCR**)^9^. For example, GRAB_eCB2.0_ sensor detects changes in 2-AG levels with a sub second temporal resolution and its activation is described in mouse neurons in primary culture and brain slices, as well as *in vivo* during behavior^10-13^. Thus, the spatiotemporal properties of such sensor provides an opportunity to study the molecular mechanism of eCB signaling at the cellular level. Here, we characterized GRAB_eCB2.0_ activation when expressed by Neuro2a (**N2a**) cells that affords the ability to further outline the pharmacological profile of its activation. We then leveraged this neuronal model system to examine stimuli-dependent increases of endogenous 2-AG production, determine their molecular mechanisms and sensitivity of ABHD6 inhibition.

## Results

### Pharmacological characterization of GRAB_eCB2.0_ expressed by N2a cells in real time

To drive GRAB_eCB2.0_ expression, we subcloned the *eCB2*.*0* DNA construct in a plasmid containing the chimeric cytomegalovirus-chicken β-actin promoter, and transfected N2a cells in culture (**Figure 1a**)^14-18^. First, we measured real time changes in fluorescence by confocal microscopy imaging (line scanning frequency: 200 Hz, 0.388 FPS) by spiking agents directly in the buffer media on the microscope’s stage (**Figure 1a**). As a positive control agonist, we selected 2-AG (i.e., the neuromodulator used to develop GRAB_eCB2.0_) formulated in buffer containing bovine serum albumin (**BSA**, 0.1 mg/ml), a lipid binding protein known to assist 2-AG’s action at CB_1_R (**Figure 1b**)^19,20^. **Figure 1c** shows that N2a cells exhibited low, yet clearly detectable, basal fluorescence at the plasma membrane, and that 2-AG (1 µM, 60 sec) increased this fluorescent signal. Similarly, both AEA (10 µM, 60 sec) and the synthetic agonist at CB_1_R, CP55940 (**CP**, 1 µM, 60 sec) increased GRAB_eCB2.0_ fluorescence at the plasma membrane (**Figure 1c**). Note that basal and agonist-triggered increase in GRAB_eCB2.0_ fluorescence reached different levels in each N2a cell as expected from a heterologous transfection approach (**Figure 1c**, see arrowhead and arrow). **Figure 1d** shows that the GRAB_eCB2.0_ protein is detected by a CB_1_R antibody by western blot, and **Figure 1e** shows that GRAB_eCB2.0_ protein expressed in N2a cells in culture is detected by a CB_1_R antibody by immunocytochemistry as well, and that its expression is heterogenous as expected for transient transfections.

**Figure 1:**
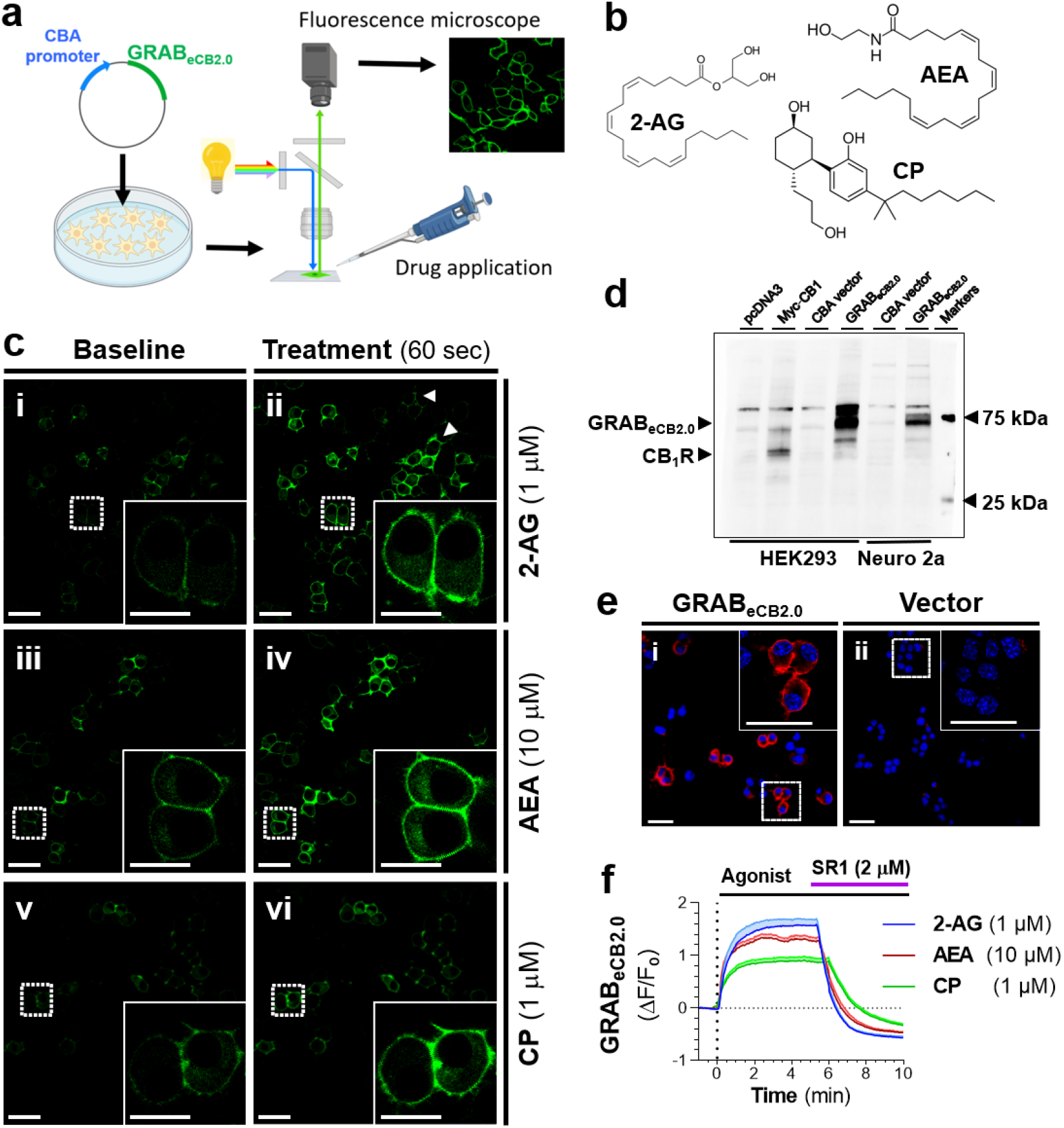
Agonist triggered increases in GRAB_eCB2.0_ fluorescence in N2a cells detected by live-cell confocal microscopy. N2a cells in culture were transfected with GRAB_eCB2.0_ plasmids and GRAB_eCB2.0_ activated by treating cells with **2-AG** (1 µM), anandamide (**AEA**, 10 µM), or CP55940 (**CP**, 1 µM), and with SR141617 (**SR1**, 2 µM). Changes in fluorescence were detected using live-cell confocal microscopy. **a]** Schematic of GRAB_eCB2.0_ plasmid transfected in N2a cells, measuring changes in fluorescence by spiking agent directly in buffer, and measuring changes in fluorescence by live-cell confocal microscopy. **b]** Chemical structures of cannabinoid agonists. **c]** Representative images of GRAB_eCB2.0_ fluorescence in N2a cells at baseline (i, iii and v: 60 sec before treatment) and 60 sec after treatment (ii, iv and vi). Heterogenous expression levels of GRAB_eCB2.0_ is illustrated with arrowhead (low expression) and arrow (high expression). Scale bars: 40 µM (insert 20 µM). **d]** Detection of GRAB_eCB2.0_ protein by western blot and detected with an antibody against CB_1_R. Positive CTR in HEK293 cells: HEK293 transfected with MYC-tagged hCB_1_R. GRAB_eCB2.0_ transfected in both HEK293 and N2a cells are reliably detected by the CB_1_R antibody. Representative western blot repeated twice with similar results. **e]** Detection of GRAB_eCB2.0_ protein express by N2a cell by ICC using an antibody against CB_1_R. N2a cells transfected with GRAB_eCB2.0_ plasmid (i) and vector only (ii). Scale bars: 20 µM (insert 20 µM). **f]** Time course of GRAB_eCB2.0_ activation by CB_1_R agonists (ΔF/F_0_), and their reversibility by SR1 applied ≈5 min after agonist treatment. N= 10-15 cells per treatment.

We then compared the time course of increased GRAB_eCB2.0_ fluorescence triggered by these CB_1_R agonists and the reduction of their responses when adding the CB_1_R antagonist SR141716 (**SR1**, 2 µM)^21^. As expected, each agonist triggered a rapid increase in GRAB_eCB2.0_ fluorescence that plateaued after approximately 60 sec, and SR1 reduced these responses within 100 sec to levels that were below the initial basal GRAB_eCB2.0_ fluorescence (**Figure 1f**). Thus, these agonists exhibited different activation profiles: 1] 2-AG and CP triggered a 2-fold steeper initial response (slope) and 2-fold greater maximal response (peak) compared to AEA; 2] 2-AG and AEA triggered a 2-fold overall greater response (area under the curve) for compared to CP; 3] their responses were blocked by SR1 treatment and led to comparable decays (*t*) in their signal (**Table 1**). These results demonstrate that GRAB_eCB2.0_ expressed by N2a cells is activated by the CB_1_R agonists 2-AG > AEA > CP and these responses are blocked by the CB_1_R antagonist SR1.

**Table 1:** Parameters of GRAB_eCB2.0_ activation by agonists and reversal by antagonism. N2a cells in culture were transfected with GRAB_eCB2.0_ plasmids and GRAB_eCB2.0_ activated by treating cells with **2-AG** (1 µM), anandamide (**AEA**, 10 µM), or CP55940 (**CP**, 1 µM), and with SR141617 (**SR1**, 2 µM). Changes in fluorescence were detected using live-cell confocal microscopy. Data are shown as a mean of 39-70 cells from 4 independent experiments; error bars represent S.E.M.

| PD parameter | Units | CB<sub>1</sub>R agonists |  |  |
| --- | --- | --- | --- | --- |
|  |  | 2-AG | CP | AEA |
| Initial response: slope | ($\times 10^{-2} \Delta F/F_0/\text{sec}$) | **3.73** | **2.30** | **4.33** |
|  | (95% CI) | (3.57-3.89) | (2.16-2.44) | (3.97-4.69) |
| Time to peak | (min) | **4.38** | **3.74** | **4.64** |
| Maximal response: peak | ($\Delta F/F_0$) | **1.79** | **0.90** | **1.69** |
|  | (std. error) | 0.074 | 0.058 | 0.150 |
| Overall response | (area under the curve) | **453** | **239** | **434** |
|  | (std. error) | 12 | 7 | 20 |
| Antagonism response: t | (sec) | **62.94** | **42.18** | **35.5** |
|  | (95% CI) | (57.71-69.08) | (40.21-44.31) | (33.22-38.03) |

### High-throughput measures of GRAB_eCB2.0_ activity in N2a cells

To define the pharmacological profile of GRAB_eCB2.0_ for use in a high throughput screening (**HTS**) format, we developed a fluorescent plate reader assay (96 wells, 3 Hz scanning frequency). Thus, the initial basal fluorescence signal of GRAB_eCB2.0_ expressed by N2a cells was measured for 1 min, agents were spiked in the media, and approximately 120 sec later, cells were reinserted in the plate reader and fluorescence was measured for an additional 30 min (**Figure 2a**). To facilitate a data analysis pipeline, we developed a MATLAB R2021a algorithm which averages the fluorescence value of each well over time, for multiple experiments and at select timepoints (see the following link for the code: https://github.com/StellaLab/StellaLab.git). **Figures 2b** and **2c** show that 2-AG induced a concentration-dependent increase in fluorescence and SR1 a concentration-dependent decrease in fluorescence. Specifically, the concentration-dependent increase in GRAB_eCB2.0_ signal was detectable at 0 sec (when cells were reinserted in the plate reader and first fluorescence signal measured), and these signals reached a peak response followed by plateau responses that could last 30 min. To calculate the EC_50_ and IC_50_ of these responses, we averaged the GRAB_eCB2.0_ signals at peak response (4-5 min for 2-AG and 11-16 min for SR1) and calculated an EC_50_ of 82 nM for 2-AG, a response that is equivalent to its potency at CB_1_R (EC_50_ = 12-100 nM)^22,23^, and an IC_50_ of 1.2 µM for SR1, which is approximately 1000-fold less than its reported K_i_ at CB_1_R (**Figures 2d**)^21^. Both AEA and CP also induced concentration-dependent increases in fluorescence with potencies of 201 nM and 190 nM, respectively, responses that are also lower than their respective activities at CB_1_R (EC_50_ = 69-276 nM and 0.05-31 nM respectively)^24,25^ (**Figures 2d** and Supplementary Figure S1). The EC_50_s values of 2-AG and AEA for GRAB_eCB2.0_ measured here are lower than their previously reported values in cell culture models systems (i.e., 2-AG = 3.1-9.0 µM and AEA = 0.3-0.8 µM)^10^, most likely explained by our addition of BSA to the buffer to facilitate solubility and interaction with CB_1_R^26^.

**Figure 2:**
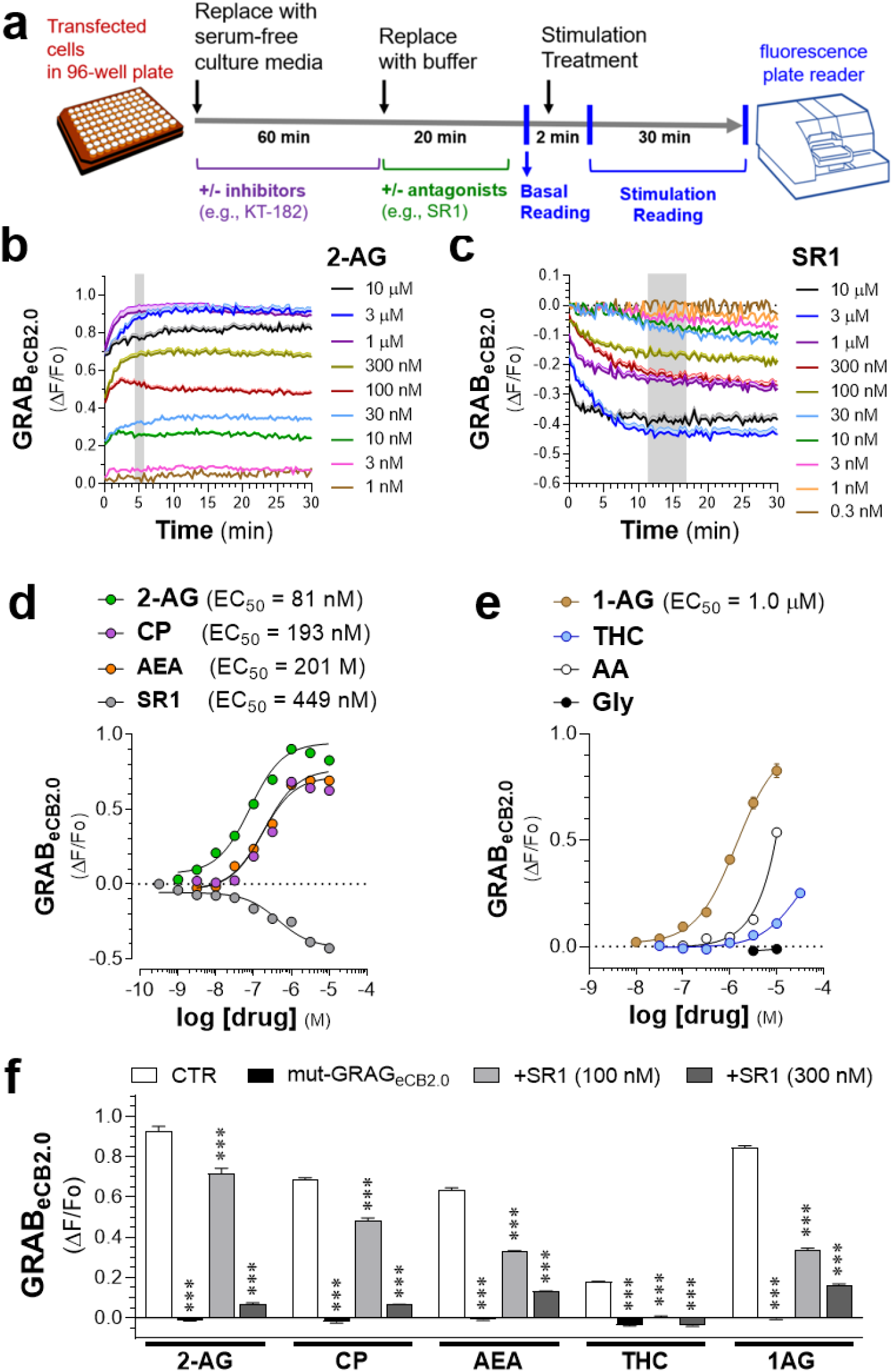
Pharmacological profile of GRAB_eCB2.0_ in N2a cells. N2a cells in culture were transfected with GRAB_eCB2.0_ plasmids and GRAB_eCB2.0_ was activated by treating cells with increasing concentrations of 2-AG, AEA, CP, 1-AG, THC, and SR1. Changes in fluorescence were detected using a 96 well plate reader. **a]** Schematic outlining protocol to measure fluorescent changes in GRAB_eCB2.0_-transfected N2a cells using a fluorescent plate reader. **b-c]** Time course of fluorescent change (ΔF/F_0_) following treatment with increasing concentrations of 2-AG or SR1. **d-e]** Concentration-dependent changes in GRAB_eCB2.0_ fluorescence triggered by 2-AG, AEA, CP, and SR1. Fluorescent signal between 4-5 min was averaged for 2-AG, AEA, CP and 1-AG, between 11-16 min for SR1 and THC, and 25-30 min for arachidonic (**AA**) and Glycerol (**Gly**) (kinetics are in Supplementary Figure S2). **f]** 2-AG (1 µM), CP (1 µM), AEA (10 µM), THC (10 µM) and 1-AG (1 µM) increases in GRAB_eCB2.0_ signal are reduced by SR1 (100 and 300 nM), and in N2a cells expressing mut-GRAB_eCB2.0_. Data are shown as mean of 3-76 independent experiments for 2-AG, 3-10 for AEA, 3-39 for CP, 3-5 for 1-AG, and 4-7 for SR1. Experiments were done in triplicate; error bars represent S.E.M.

We next tested the 2-AG analog, 1-arachidonoylglycerol (**1-AG**), and found that it increased GRAB_eCB2.0_ fluorescence with an EC_50_ of 1 µM and is comparable to its 1-2 µM potency at wild-type CB_1_R (**Figures 2e** and Supplementary Figure S1)^22,27^. Glycerol, a hydrolysis product of 2-AG and 1-AG, did not influenced GRAB_eCB2.0_ signal at 3 and 10 µM, whereas arachidonic acid (**AA**), the other hydrolysis product of 2-AG and 1-AG, significantly increased GRAB_eCB2.0_ signal at 3 and 10 µM (**Figures 2e** and Supplementary Figure S1).

To further characterize the pharmacological profile of GRAB_eCB2.0,_ we examined the pharmacodynamic response of the phytocannabinoid Δ^9^-tetrahydrocannabinol (**THC**), which induced a small, concentration-dependent increase in fluorescence, as expected for a partial agonist at CB_1_R, but with an EC_50_ of 1.3 µM, which is 10-100-fold less potent than its activity at CB_1_R^28^ (**Figure 2e** and Supplementary Figure S1). Pre-treatment of N2a cells with SR1 followed by application of 2-AG, AEA, CP, 1-AG, and THC blocked activation of GRAB_eCB2.0_, and these agonists also failed to elicit increases in fluorescence when transfecting mutant GRAB_eCB2.0_ (**mut-GRAB**_**eCB2.0**_), in which phenylalanine 177 has been mutated to alanine in the region within the orthosteric binding pocket of the CB_1_R^29,30^ (**Figure 2f**). Together, these results show that changes in GRAB_eCB2.0_ fluorescence are reproducibly detected using a 96 well plate-reader, that 2-AG and 1-AG increase this signal with EC_50_s comparable to their activities at the CB_1_R, and that GRAB_eCB2.0_ detects AEA, CP, THC and AA although at 10-100-fold lesser potency than CB_1_R.

### G protein-coupled receptors versus ionotropic stimuli induce differential increases in 2-AG

N2a cells express metabotropic G protein-coupled receptors (**GPCRs**) that couple to G_q_ proteins, phospholipase C (**PLC**) and DAGL, as well as calcium-permeable ionotropic receptors, all molecular components involved in increasing 2-AG production^18,31^,32. We selected bradykinin (**BK**) to stimulate metabotropic receptor-dependent production of 2-AG^33^, and adenosine triphosphate (**ATP**) to stimulate ionotropic receptor-dependent production of 2-AG, and measured changes in GRAB_eCB2.0_ signal by live cell fluorescence microscopy (**Figure 3a**)^34,35^. We also tested ionomycin, a receptor-independent calcium ionophore that increase _i_[Ca^2+^] and ensuing 2-AG production in neurons, and mastoparan, a direct activator of Gq proteins^22,36^,37. **Figure 3b** shows that BK (1 µM) caused a transient increase in GRAB_eCB2.0_ fluorescence that reached its maximal response within 2-3 min, and that this response decayed by approximately half after 10 min. Using LC-MS as comparison readout, we found that BK increased 2-AG levels by 94% after 2 and 72% after 10 min (**Figure 3c**; LC-MS detection limits are in Supplementary Figure S3a). **Figure 3d** shows that ATP (300 µM) also triggered a pronounced and transient increase in GRAB_eCB2.0_ fluorescence that reached a maximum of 0.38-0.4 ΔF/F between 7-9 min. ATP also significantly increased 2-AG levels by 53% after 2 min when measured by LC-MS, but remarkably this response returned to baseline after 10 min, in sharp contrast to the ATP-induced GRAB_eCB2.0_ response which remained elevated after 10 min treatment (**Figure 3d** and **3e**). Combining BK (1 µM) and ATP (300 µM) produced a strong and prolonged increase in GRAB_eCB2.0_ fluorescence that reached a maximum of 0.56-0.58 ΔF/F that lasted between 3-6 min, which decayed by only 20% after 10 min (**Figure 3f**). This response was paralleled by 129% and 67% increases in LC-MS 2-AG levels after 2 and 10 min, respectively (**Figure 3g**). AEA remained below detection limit under basal and stimulated conditions as quantified by LC-MS, indicating that the main eCB produced under these conditions is 2-AG (Supplementary Figure S3a).

**Figure 3:**
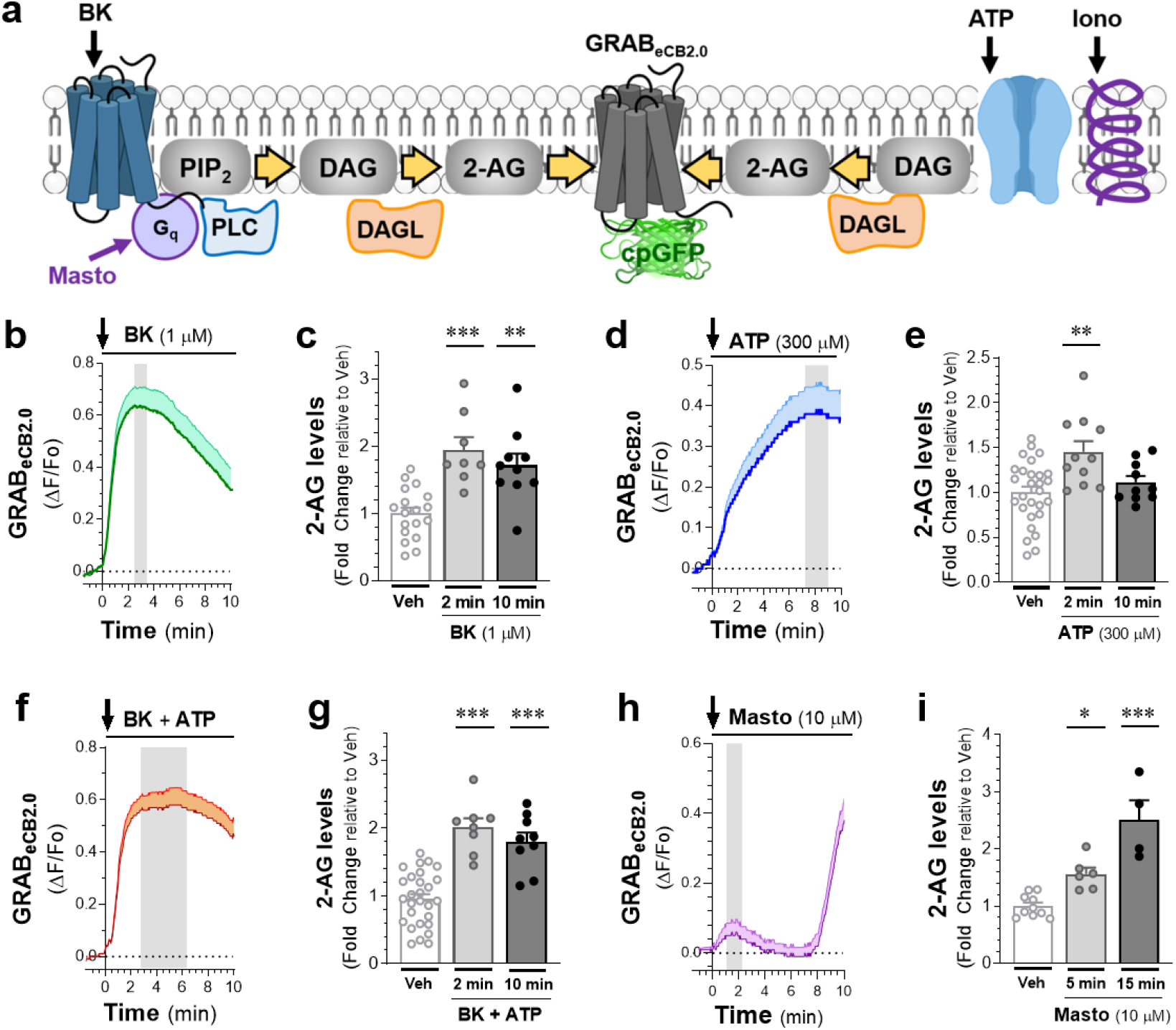
Stimuli-dependent increase in 2-AG levels in N2a cells. N2a cells were transfected with GRAB_eCB2.0_ and activation of GRAB_eCB2.0_ tested by treating cells with bradykinin (**BK**), adenosine triphosphate (**ATP**) and Mastoparan (**Masto**), and fluorescent changes detected by live cell microscopy and liquid chromatography mass spectrometry (**LC-MS**). **a]** Illustration of 2-AG production regulated by distinct receptor-dependent (using ATP and bradykinin) and receptor-independent (using ionomycin (**Iono**) and Masto) mechanisms. **b-c]** BK (1 µM) response on GRAB_eCB2.0_ fluorescence as measured by live-cell imaging (b) and 2-AG levels as measured by LC-MS after 2 and 10 min of treatment (c). **d-e]** ATP (300 µM) response on GRAB_eCB2.0_ fluorescence as measured by live-cell imaging (d) and on 2-AG levels as measured by LC-MS at 2 and 10 min of treatment (e). **f-g**] BK (1 µM) + ATP (300 µM) response on GRAB_eCB2.0_ fluorescence as measured by live-cell imaging (f) and on 2-AG levels as measured by LC-MS at 2 and 10 min of treatment (g). **h-i]** Masto (10 µM) response on GRAB_eCB2.0_ fluorescence as measured by live-cell imaging (h) and on 2-AG levels as measured by LC-MS at 2 and 15 min of treatment (i). Data is shown as a mean of 39-51 cells from 3-4 independent experiments for live cell imaging and N= 4-18 independent experiments for LC-MS; error bars represent S.E.M.

We then tested receptor-independent stimuli, mastoparan and ionomycin, and found smaller and more transient increases in GRAB_eCB2.0_ signal. Specifically, mastoparan (1 µM) triggered a small and slow increase in GRAB_eCB2.0_ signal that reached a maximum of 0.06-0.07 ΔF/F, this response started to plateau at 2 min, returned to baseline by 6 min, and remarkably rose again after 8 min (**Figure 3J**). This response was not paralleled by the increased 2-AG levels as measured by LC-MS (increases by 57% after 5 min and 150% after 15 min of treatment) (**Figure 3K**). Mastoparan is known to trigger the formation and release of membrane vesicles after 10 min of treatment and accordingly we delectated such vesicles that also contained GRAB_eCB2.0_ (Supplementary figure S3a). Ionomycin (2.5 µM) also triggered a small and slow increase in GRAB_eCB2.0_ signal that reached a maximum of 0.06-0.07 ΔF/F, this response started to peak at 6 min and returned to basal by 10 min (Supplementary figure S3b). Surprisingly, when measured by LC-MS, ionomycin increased 2-AG levels by 54% after both 2 min and 10 min of treatment, suggesting that the GRAB_eCB2.0_ fluorescence at the plasma membrane may not detect intracellular 2-AG (Supplementary figure S3b). Together, these results show that GRAB_eCB2.0_ reliably detects changes in 2-AG levels induced by distinct stimuli and occurring with differing kinetics. Increases in 2-AG levels by these stimuli were also detected in parallel by LC-MS, although with a more variable sensitivity as compared to the GRAB_eCB2.0_ signal.

### Isolating the mechanisms underlying the BK-, ATP- and G protein-mediated increases in 2-AG

To identify the receptors and lipases that mediate these stimuli-dependent increases in 2-AG levels, we leveraged selective antagonists and inhibitors and measured changes in GRAB_eCB2.0_ fluorescence using our HTS-N2a model system (**Figure 4a**). **Figure 4b** shows that BK triggered a concentration-dependent increase in GRAB_eCB2.0_ signal characterized by a peak response at 4-5 min, a maximal response reached at 3-10 µM, a progressive acceleration of GRAB_eCB2.0_ activation (slope between 1-3 min) and diminishing decay of this response (5-30 min). Considering the exquisite sensitivity of the GRAB_eCB2.0_ signal to the concentration-dependent increase in 2-AG, we translated the BK (1 µM)-stimulated increase in GRAB_eCB2.0_ signals to 2-AG concentrations and estimated that BK (1 µM) increased 2-AG levels by ≈ 8 nM (Supplementary Figure S4). Mechanistically, the BK (1 µM) response measured between 4-5 min was, as expected, blocked by SR1 (300 nM) and absent in N2a cells transfected with mut-GRAB_eCB2.0_ (**Figure 4c**). Furthermore, the BK response was blocked by the selective B_2_R antagonist, HOE140 (300 nM)^38^, indicating the involvement of B_2_R metabotropic receptors expressed by N2a cells^39-41^ (**Figure 4c**). The BK response was also blocked by a selective DAGL inhibitor (**DO34**, 10 nM) (**Figure 4c**), a concentration that fully inhibits DAGL activity in N2a as measured by activity-based protein profiling (**ABPP**, Supplementary Figure S4)^5,42^. To test the calcium-dependence of this response, we increased intracellular calcium levels by co-treating BK (1 µM) with either ionomycin (2.5 µM) or thapsigargin (**TG**, 10 µM), a Sarco/endoplasmic reticulum calcium-ATPase (**SERCA ATPase**) inhibitor. To test the effect of decreased calcium, we either removed calcium from the buffer or removed calcium from the buffer and added EGTA (1 mM). We found that the BK response was not affected by ionomycin, unchanged by removing _e_[Ca^2+^], slightly decreased by the absence of _e_[Ca^2+^] and was increased by 2-fold by TG (**Figure 4c**). Importantly, these treatments did not affect the GRAB_eCB2.0_ signal when introduced as single agents or in combination with 2-AG (1 µM), emphasizing their selectivity (Supplementary Figure S4). Both absence of extracellular calcium and TG had a small reducing effect on basal GRAB_eCB2.0_ signal after 25-30 min incubation (Supplementary Figure S4). Thus, BK increased 2-AG production through the activation of B_2_K receptors that couple to DAGL, and this response is enhanced by increases in _i_[Ca^2+^].

**Figure 4:**
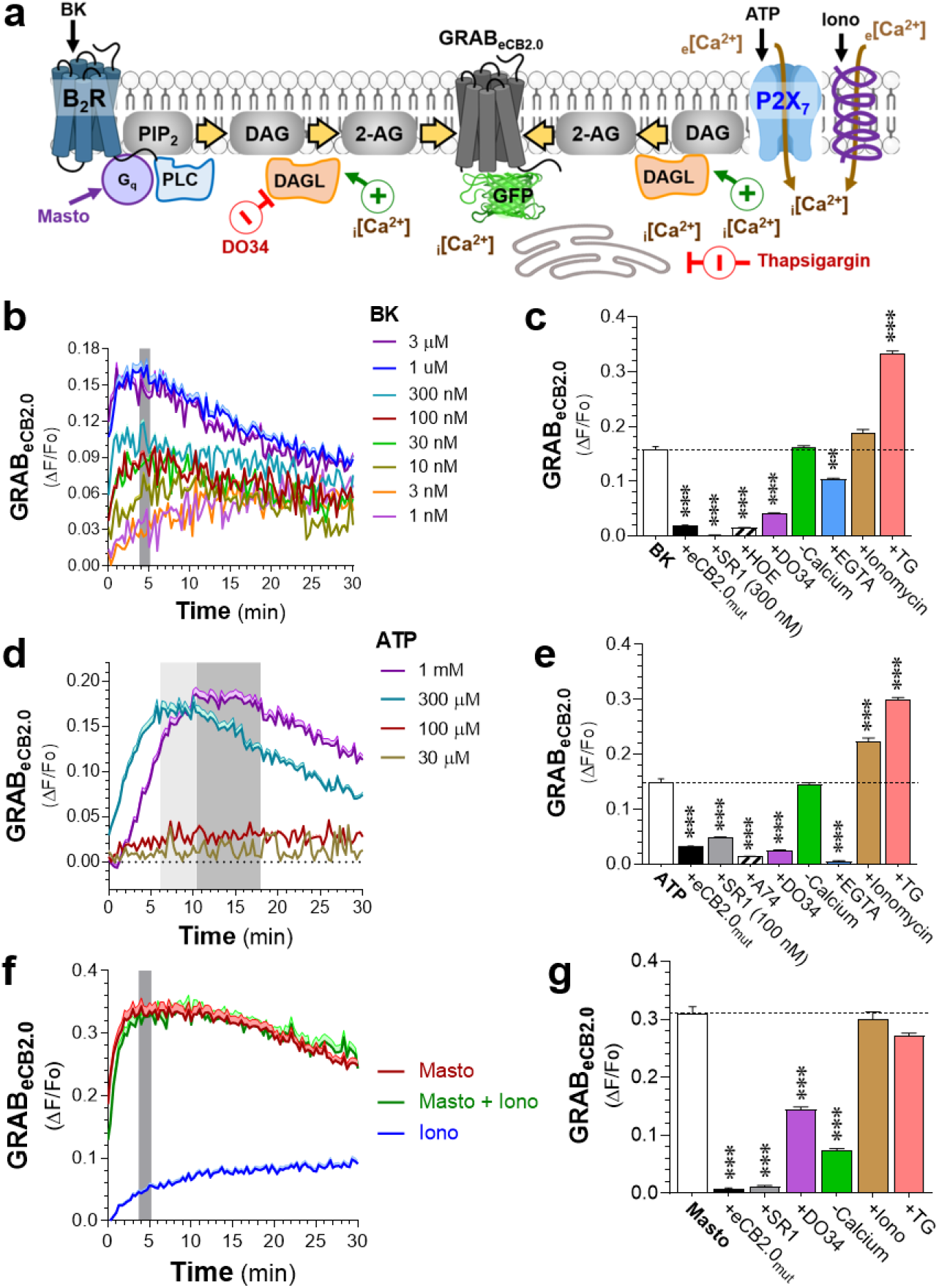
Mechanisms that mediate the BK-, ATP- and mastoparan-triggered increase in 2-AG levels in N2a cells. N2a cells were transfected with GRAB_eCB2.0,_ cells were treated with bradykinin (**BK**), adenosine triphosphate (**ATP**), ionomycin (**Iono**) and Mastoparan (**Masto**), and changes in GRAB_eCB2.0_ fluorescence detected with a 96 well plate reader. **A]** Mechanism of 2-AG production was tested by using D034 (10 nM) to inhibit DAGL. Calcium-dependence of this effect was tested by increasing intracellular calcium _i_[Ca^2+^] using thapsigargin (**TG**, 10 µM) and Iono (2.5 µM), and absence of calcium (-Calcium) and chelating calcium (**EGTA**, 1 mM). **b-c]** Concentration-dependent effect of BK (b) and ATP (c) on GRAB_eCB2.0_ fluorescence. Peak response reached after 4-5 min for BK, and 6-11 min for ATP (300 µM, light gray) and 9-18 min or ATP (1 mM, dark gray). N = independent experiments: 3-21 for BK and 4-23 for ATP. **d-e]** Mechanism that mediates BK and ATP responses. N2a cells expressing GRAB_eCB2.0_ or mut-GRAB_eCB2.0_ were treated with BK (1 µM, C) and ATP (300 µM, D), as well as either HOE 140 (**HOE**, 1 µM), A740003 (**A74**, 10 µM), D034 (10 nM), Iono (2.5 µM), or TG (10 µM). N=3-11 independent experiments; error bars represent SEM. **f]** Time course of increase in GRAB_eCB2.0_ fluorescence when treating N2a cells with Masto, Iono, or their combination. Data is shown as mean of 8-32 independent experiments; error bars represent S.E.M. **g]** Mechanism that mediates Masto responses (similar treatment as in d-e). N=3-11 independent experiments; error bars represent S.E.M.

ATP treatment however, initiated a contrasting increase in GRAB_eCB2.0_ fluorescence characterized by a maximal response of 0.18-0.2 ΔF (i.e., an increase in ≈9 nM 2-AG) that reached its maximum between 6-11 min with 300 µM, and between 9-18 min with 1 mM (**Figure 4d**). By contrast to the BK response, the ATP 300 µM- and 1 mM-increased GRAB_eCB2.0_ signal had comparable initial activation slopes (between 2-5 min) and similar decays following their plateau response (**Figure 4d** and Supplementary Figure S4). Thus, we sought to investigate the mechanism at ATP 300 µM. **Figure 4e** shows that the ATP (300 µM)-triggered response measured between 6-11 min which was antagonized by SR1 (300 nM) and absent in N2a cells transfected with mut-GRAB_eCB2.0_ sensor. **Figure 4e** also shows that this response was blocked by the P2X_7_ antagonist, A74003 (10 µM)^43^, indicating that it is mediated through this nonselective ligand-gated cation channel that is only activated by high micromolar – millimolar concentrations of ATP^44^, is expressed by N2a cells^45^, and controls 2-AG production in microglia^35^. The ATP (300 µM)-triggered GRAB_eCB2.0_ signal was blocked by DO34 (10 nM), unaffected when removing extracellular calcium, completely blocked in the absence of _e_[Ca^2+^] and increased by both ionomycin and TG (**Figure 4e**). Thus, ATP (300 µM) triggers 2-AG production through P2X_7_ receptors that activate DAGL, and this response is greatly sensitive to both _i_[Ca^2+^] and _e_[Ca^2+^].

Ionomycin increased GRAB_eCB2.0_ fluorescence starting immediately after treatment, and this response reached linearity after 10 min and a 0.09 ΔF response after 30 min (**Figure 4f**). Mastoparan triggered a rapid and pronounced increase in GRAB_eCB2.0_ signal that reached its peak response at 4-5 min and slowly decayed by ≈20% after 30 min (**Figure 4f**) This response was not affected by adding ionomycin (**Figure 4f**). **Figure 4g** shows that the mastoparan response measured between 4-5 min was: blocked by SR1 (300 nM) and absent in mut-GRAB_eCB2.0_-transfected N2a cells, greatly reduced by both D034 (10 nM) and the absence of calcium in the buffer and was unaffected by ionomycin and TG (**Figure 4g**). Taken together, these results indicate that BK, ATP, ionomycin and mastoparan increases 2-AG levels via discrete mechanisms and different calcium-dependencies.

### ABHD6 selectively controls metabotropic-dependent increase in 2-AG

To study if and how ABHD6 inhibition influences B_2_K- and P2X_7_-stimulated increase in GRAB_eCB2.0_ signal in N2a cells, we first measured in situ ABHD6 activity and determined its sensitivity to the ABHD6 inhibitor, KT-182^46^. As previously reported, ABPP analysis of N2a cells confirmed pronounced endogenous ABHD6 activity and absence of MAGL activity^47-49^ (**Figure 5a**). This activity was inhibited by KT-182 (IC_50_ of 3.4 nM); and KT-182 (10 nM) inhibited 96% of ABHD6 activity while not affecting the activity of multiple serine hydrolases, including FAAH (**Figures 5b** and **5c**). Using the HTS plate reader approach, we found that KT-182 (10 nM) enhanced the 2-AG (1 µM)-triggered increase in GRAB_eCB2.0_ signal by ≈50% without affecting the overall kinetic of this response, and without affecting the CP-triggered increase in GRAB_eCB2.0_ signal, emphasizing the selectivity of KT-182 (10 nM) inhibition of endogenous ABHD6 activity (**Figure 5d** and **5e**). **Figure 5f** and **5g** show that KT-182 (10 nM) enhanced both the BK (1 µM)- and the mastoparan (1 µM)-triggered increase in GRAB_eCB2.0_ fluorescence. By sharp contrast, the ATP (300 µM)-triggered increase in GRAB_eCB2.0_ fluorescence was not affected by KT-182 (10 nM) (**Figure 5h**). **Figure 5i** shows that the increased GRAB_eCB2.0_ signal triggered by the combination of BK (1 µM) + ATP (300 µM) was enhance by KT-182 (10 nM), indicating that the metabotropic component of this stimuli is enhanced when ABHD6 is inhibited. Together, these results suggest that ABHD6 preferentially controls metabotropic-dependent increased 2-AG production over ionotropic-dependent increased 2-AG production.

**Figure 5:**
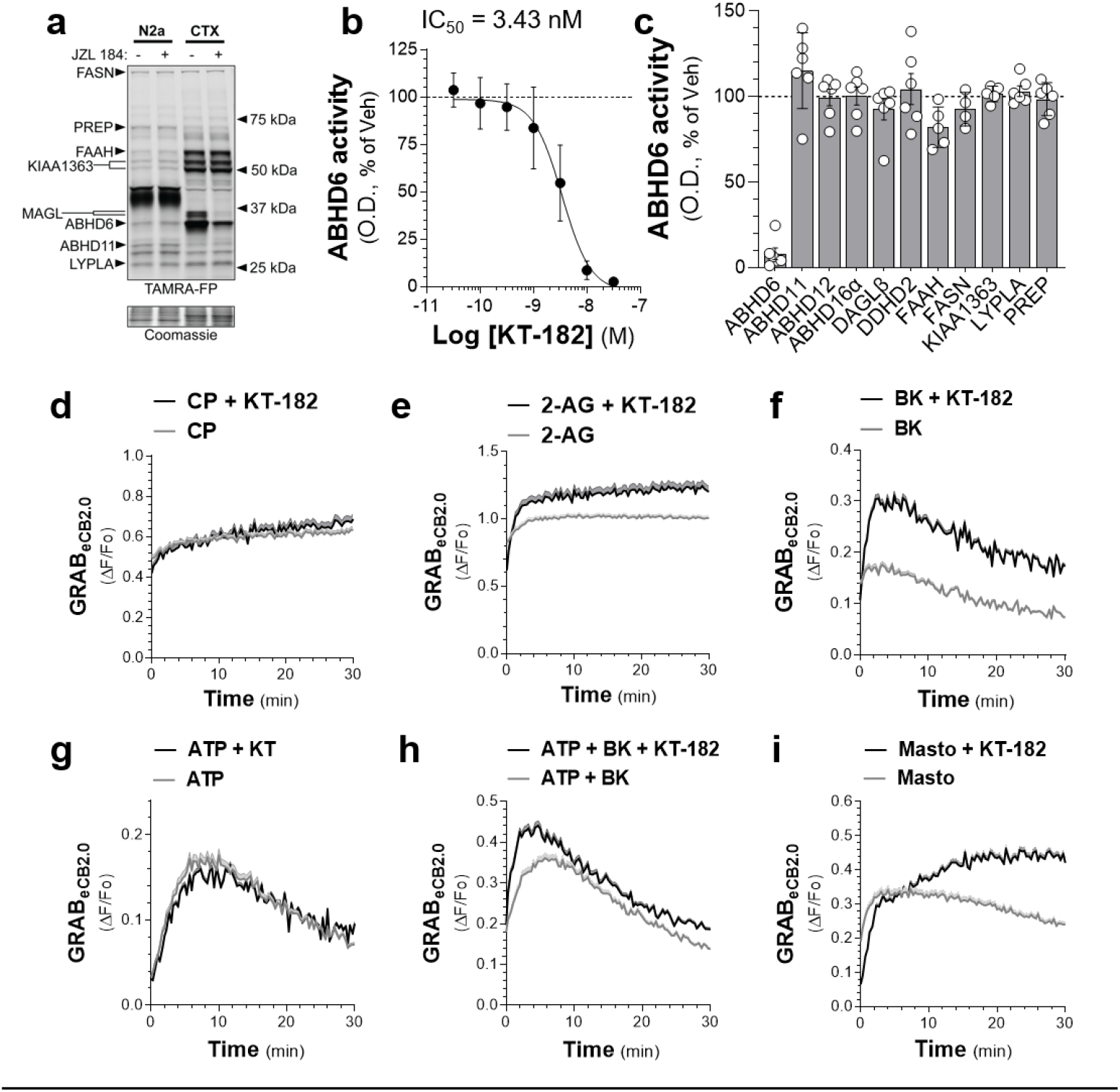
ABHD6 activity and inhibition in N2a cells and its selective control of metabotropic-dependent increase in 2-AG levels. ABHD6 and related enzymatic activities in N2a cells were measure by ABPP. N2a cells were transiently with GRAB_eCB2.0_ and activation of GRAB_eCB2.0_, cells treated with 2-AG, CP, bradykinin (**BK**), adenosine triphosphate (**ATP**), ionomycin (**Iono**), mastoparan (**Masto**) and KT-182 (10 nM), and changes in fluorescence detected with a 96 well plate reader. **a]** Absence of MAGL activity as shown by using gel based ABPP of N2a cells proteome. Positive control: mouse brain cortex membrane proteome (**CTX**). Membrane proteome was treated with vehicle or JZL 184 (1 µM, 30 min) followed by the TAMRA-FP activity-based probe (500 nM, 15 min). **b]** KT-182 inhibition of endogenous ABHD6 activity in intact N2a cells in culture treated for 60 min. ABHD6 activity was measured using gel-based ABPP with the MB064 activity-based probe (2 µM, 15 min). **c]** KT-182 (10 nM, 60 min in situ) selectively inhibits ABHD6 over other related enzymes in N2a cells. (b-c: N = 3-4 independent experiments and error bars represent S.E.M.) **d-i]** Effect of KT-182 (10 nM, added during the 60 min serum-free cell culture preincubation) on stimulated increase in GRAB_eCB2.0_ fluorescence in N2a cells. N2a cells were pretreated with KT-182 (10 nM) and treated with CP (1 µM, D), 2-AG (1 µM, E), BK (1 µM, F), ATP (300 µM, G), BK + ATP (1 µM & 300 µM, H), and Masto (10 µM, I). Data are shown as mean of 3-16 independent experiments; error bars represent S.E.M.

## Discussion

Here we report the pharmacological profile of GRAB_eCB2.0_ sensor expressed by neuronal cells in culture and found that it has an exquisite sensitivity to changes in 2-AG concentrations, and that it also detects AEA, CP, THC and AA although at 10-100-fold lesser potency than CB_1_R. Leveraging this model system, we isolated the mechanisms that mediate the BK-, ATP- and mastoparan-stimulated increase in 2-AG production in N2a cells, and discovered that ABHD6 preferentially controls metabotropic-dependent increases in 2-AG production over ionotropic-dependent increases in 2-AG production.

Our in vitro characterization of GRAB_eCB2.0_ demonstrates that this sensor is activated by 2-AG with comparable potency as CB_1_R, and it is 100-1000-fold less sensitive to AEA, CP, and SR1. Thus, stimuli-dependent increases in GRAB_eCB2.0_ fluorescence is unlikely to report changes in AEA levels as this eCB is typically 10-100-fold less abundant in cells compared to 2-AG, and GRAB_eCB2.0_ is only activated by AEA at high nanomolar concentrations. Of note, although exhibiting a lower affinity to SR1 than the wild-type CB_1_R, the stimuli-dependent increase in GRAB_eCB2.0_ fluorescence could be competitively blocked by this CB_1_R antagonist, but only at high 100-300 nM concentrations. Furthermore, the GRAB_eCB2.0_ sensor is expressed by the entire plasma membrane where it detects discrete changes in 2-AG within seconds that cannot be captured by LC-MS that measures whole cell 2-AG content. The field if endocannabinoid research has relied primarily on mass spectrometry-based approaches to quantify eCB levels in biological matrices, an important body of work that defined their amounts in select brain areas and cells types in culture, and unraveled the pronounced changes in eCB tone associated with physiopathological states^50-52^. However, limitations to these techniques include variable eCB recovery due to loss of eCBs during sample preparation, low throughput of sample analysis and limited spatiotemporal resolution due to tissue sampling. The recent development of GRAB_eCB2.0_ enabled the precise measurements of nanomolar changes within seconds and in a cell specific manner. Accordingly, the BK and ATP stimulated increase in 2-AG levels in N2a cells is detected by LC-MS analysis, however, with a notable lower temporal resolution and magnitude, as well as pronounced experimental variability. By contrast, the BK and ATP triggered increase in GRAB_eCB2.0_ fluorescence is detected within sec of treatment and with low variability between experiments. The disparity between the results from mass spectrometry and the GRAB_eCB2.0_ sensor in the case of ionomycin and 300 µM ATP may reflect the higher spatial and temporal resolution of this sensor compared to LC-MS. Thus, the discrepancy in the GRAB_eCB2.0_ signal and LC-MS results may reflect that LC-MS is detecting total 2-AG content in entire cells (plasma membranes and intracellular organelles), while the GRAB_eCB2.0_ signal is likely detecting 2-AG changes localized to the plasma membrane. To our knowledge, our study is the first to directly compare between change in 2-AG by LC-MC and GRAB_eCB2.0._ fluorescence in the same model system.

2-AG production can be increased by distinct stimuli and molecular mechanisms. For example, GPCR dependent activation of Gq protein and increased PLC activity produces DAG, and thus enhances substrate availability for DAGLα and β, whose activities are potentiated by increases in _i_[Ca^2+^]^31,34,53-57^. Here we show that B_2_R-triggered increase in GRAB_eCB2.0_ signal is potentiated by TG that blocks SERCA channels and increases _i_[Ca^2+^]. By contrast, the P2X_7_-mediated increase in GRAB_eCB2.0_ signal is not affected by TG and dependent on _e_[Ca^2+^] as expected for a calcium-permeable ligand-gated ion channel. One possible explanation for this differential control by ABHD6 of stimuli-dependent increases in 2-AG levels is that this enzyme might be in closer proximity to B_2_K receptors compared to P2X_7_ receptors, which would suggest that the involvement of ABHD6 in the control of eCB signaling might depend on its subcellular expression pattern. Initial evidence indicates that ABHD6 only regulates 2-AG levels and its activity at CB_1_R in neurons stimulated by neurotransmitters^4^. Accordingly, KT-182 has no effect on basal GRAB_eCB2.0_ signal and the CP-stimulated GRAB_eCB2.0_ signal. The preferential ABHD6-dependent enhancement of BK-stimulate increases in 2-AG production compared to ATP-stimulated increases in 2-AG production suggest subcellular compartmentalization of ABHD6 to intracellular domains where metabotropic Gq-coupled receptors reside. It should be emphasized that we used non-differentiated N2a cells in culture as model system and thus changes in GRAB_eCB2.0_ signal are likely to reflect 2-AG autocrine signaling. Accordingly, our results combined with published studies on 2-AG-CB_1_R signaling suggest at least four scenarios: post-synaptic production of 2-AG can activate: 1) presynaptic CB_1_R, 2) neighboring CB_1_R expressed by glia, 3) CB_1_R expressed by mitochondria and, our results, 4) autocrine CB_1_Rs.

The B_2_K-dependent increase in 2-AG levels can be significantly enhanced when inhibiting ABHD6, suggesting that such an inhibitor may have therapeutic value for the treatment of neurological diseases involving BK and CB_1_R, such as chronic pain. Considering that ABHD6 inhibitors exhibit promising therapeutic efficacy in many preclinical mouse models of neurological diseases; including models of seizures, traumatic brain injury, and experimental autoimmune encephalomyelitis^58-61^, our results might help better understand the therapeutic MOA of ABHD6 inhibitors, optimize their efficacy, as well as help identify therapeutic indications. Importantly, the therapeutic efficacy of ABHD6 inhibitors does not appear to undergo tolerance or trigger overt side-effects, two positive qualities for potential small molecule therapeutics^61,62^.

In conclusion, we developed and characterized a cell culture model system with HTS capacity for examining in depth the mechanisms of stimuli-dependent changes in 2-AG production. We established a 2-AG detection using the newly developed GRAB_eCB2.0_ sensor within seconds and discovered that ABHD6 preferentially controls metabotropic-dependent increase in 2-AG production over ionotropic-dependent increase in 2-AG production. Thus, our study shows that ABHD6 selectively controls stimuli-dependent increases in 2-AG production and emphasizes its specific role in eCB signaling and the potential of ABHD6 inhibitors as novel therapeutics for the treatment of neurological diseases.

## Acknowledgements

This work was supported by the National Institutes of Health (NS118130 and DA047626 to N.S., DA055448 to AE, DA033396 to MRB, and T32GM007750). We would also like to acknowledge support from the University of Washington Center of Excellence in Opioid Addiction Research/ Molecular Genetics Resource Core (P30DA048736). National Natural Science Foundation of China (31925017 and 31871087), the Beijing Municipal Science & Technology Commission (Z181100001318002 and Z181100001518004), the NIH BRAIN Initiative (1U01NS113358), the Shenzhen-Hong Kong Institute of Brain Science (NYKFKT2019013), the Science Fund for Creative Research Groups of the National Natural Science Foundation of China (81821092) and grants from the Peking-Tsinghua Center for Life Sciences and the State Key Laboratory of Membrane Biology at Peking University School of Life Sciences (to YL).

## Methods

### Chemicals and Reagents

Ionomycin (Tocris, 1704), mastoparan (Sigma Aldrich, M-5280), bradykinin (Sigma Aldrich), adenosine triphosphate, thapsigargin (Tocris, 1138), HOE 140 (Tocris, 3014), A740003 (Tocris, 3701), 2-arachidonoylglycerol (Cayman), 1-arachodonoylglycerol (Cayman), THC, CP55940, SR141716, arachidonoylethanolamide (Cayman), D034, KT-182, goat anti-CB_1_R antibody (1:1000 for IF and 1:2,500 for immunoblotting); AlexaFluor 647 conjugated donkey anti-goat (1:1000); and rabbit anti-actin (1:2,500; Sigma Aldrich); IRDye 800 CW conjugates donkey anti-goat (LI-COR); IRDye 680 RD conjugated goat anti-rabbit (LI-COR).

### Cloning

GRAB_eCB2.0_ and mut-GRAB_eCB2.0_ DNA were subcloned into an AM/CBA-WPRE-bGH plasmid using the BamHI and EcoRI restriction sites. The plasmid was purified (Purelink HiPure Plasmid Maxiprep Kit, Invitrogen, CA) from transformed Stellar Competent Cells (Takara Bio Inc, Japan). The DNA was sequenced and verified (CLC Sequence Viewer 8) prior to use in transfection.

### Cell Culture

Neuro2a cells (gift from Dr. John Scott) were grown in DMEM (supplemented with 10% fetal bovine serum and 1% penicillin/streptomycin) at 37°C and 5% CO_2_. To passage cells for experiments, a confluent 10 cm plate of cells was detached by incubating with 0.25% Trypsin-EDTA for 2-3 minutes at 37°C, adding 4-5 mls of supplemented DMEM and using gentle pipetting to remove any cells still attached, then added to a new plate with fresh supplemented DMEM. Cells were passaged every 3-4 days, and for no more than 25 passages.

### Transfection

All transfections were done by incubating DNA with the transfection reagent polyethylenimine (PEI, 25K linear, Polysciences 23966) in a 1:3 ratio in serum free DMEM, incubating for 20-30 min. The DNA/PEI mixture was then added to cells in a dropwise fashion (or directly into media in eCB2.0 96 well-plate assay) without changing the growth media. Cells were transfected when they were at least 50% confluent and were incubated for 24 hours after transfection before harvesting or using for eCB2.0 assays.

### Immunofluorescence

Glass coverslips (Fisher Scientific, 12-545-82) contained in a 6-well plate were coated with poly-D-lysine (50 ng/ml, Sigma, P6407) for 1-2 h at 37°C after which the poly-D-lysine was removed, and coverslips were washed 3 times with sterile water and one time with DMEM. N2a cells were detached and resuspended in supplemented DMEM as described above, counted using a hemocytometer, plated at a density of 100,000 cells/well, and were transfected after 24 hours with 0.75 µg DNA. 24 hours after transfection, media was removed and cells were fixed with 4% paraformaldehyde in PBS (Alfa Aeser) for 20 min at room temperature. Following fixation, the cells were washed five times with PBS and permeabilized and blocked with 0.1% saponin made and 1% bovine serum albumin made in PBS for 30 min at room temperature. Cells were incubated in goat anti-CB_1_R antibody (1:1000) overnight at 4°C. Cells were then washed with PBS 6X and incubated in AlexaFluor647 conjugated donkey anti-goat secondary antibody (Invitrogen, 1:1000) for 1 hour at room temperature. Cells were washed with PBS 6X, air dried overnight, and mounted using ProLong Diamond Antifade Mountant with DAPI (ThermoFisher, P36966). Cells were imaged with a LeicaSP8X confocal microscope using a 40X oil objective.

### Western Blotting

N2a cells were plated as described above at a density of 500,000 cells/well in a 6-well plate 24 hours after plating and were transfected the following day with 0.75 µg DNA. 24 hours after transfection, media was removed, cells were washed three times with ice cold PBS, and in the last wash cells were harvested with a cell scraper and pelleted by centrifuging at 500 x g for 10 minutes. The supernatant was aspirated, and cell pellets were kept at -80°C until further use. To make cell lysate, cells were thawed on ice, resuspended in lysis buffer (25 mM HEPES pH 7.4, 1 mM EDTA, 6 mM MgCl_2_, and 0.5% CHAPS), Dounce homogenized on ice (20-30 strokes), and incubated on a rotator at 4°C for one hour. Lysate was then centrifuged at 700 x g for 10 min at 4°C, supernatant was collected, and protein concentration of supernatant was determined using a DC Protein Assay. Samples were then mixed with 4X Laemmli Sample Buffer containing 10% β-mercaptoethanol and incubated at 65°C for 5 min. 25 µg of protein were loaded onto a 10% polyacrylamide gel and transferred to PVDF membrane. After transfer, membrane was washed once with tris-buffered saline (TBS) and incubated in blocking buffer (5% BSA in TBS) for 1 hour at room temperature, followed by incubation in primary antibody overnight at 4°C. After incubation in primary antibody, membranes were washed with TBS with 0.05% TWEEN-20 (TBST) 3 times, 10 min each. Membranes were then incubated in secondary antibody (1:10,000) for 1 hour at room temperature. Blots were then washed with TBST 3 times, 10 min each followed by 3 washes with TBS, 10 min each. Fluorescent signal was detected using a Chemidoc MP (Biorad). Primary antibodies were diluted in blocking buffer. Secondary antibodies were diluted in 1:1 TBS:Odyssey blocking buffer (LI-COR, 927-50000) Primary antibodies: goat anti-CB_1_R primary (1:2,500) and rabbit anti-actin (1:2,500; Sigma Aldrich). Secondary antibodies: IRDye 800 CW conjugates donkey anti-goat (Licor) and IRDye 680 RD conjugated goat anti-rabbit (Licor).

### Isolating N2a Membrane Proteome

Cells were washed twice by adding ice cold dulbecco’s phosphate buffered saline to attached cells and aspirating. Then cells were harvested by adding 1-2 more milliliters of ice cold dPBS and detaching cells with a cell scraper. Cells were pelleted by centrifuging at 500 x g for 10 minutes. The supernatant was discarded and pellet was stored at -80°C until further use. To isolate the crude membrane fraction of the cells, the cell pellets were thawed on ice, resuspended in lysis buffer (20 mM Hepes pH 7.2, 2 mM DTT, 10 U/mL Benzonase), dounce homogenized with 20-30 strokes, and centrifuged at 100,000 x g for 45 minutes (Beckman coulter rotor Ti55). The supernatant was discarded, and the pellet was resuspended in buffer (20 mM HEPES pH 7.2 and 2 mM DTT). Protein concentration of sample was determined (DC protein assay, Biorad) before aliquoting and flash freezing samples in liquid nitrogen. Samples were stored at -80 until use.

### Live-cell imaging

Glass bottom cell culture plates (MatTek, P35G-1.5-14-C) were coated with poly-D-lysine (50 ng/ml, Sigma, P6407) for 1-2 h at 37°C, after which the poly-D-lysine was removed, and coverslips were washed 3 times with sterile water and one time with DMEM. N2a cells were detached and resuspended in supplemented DMEM as described above, counted using a hemocytometer, plated (250,000 cells per well) and were transfected after 24 hours with 0.75 µg DNA. 24 hours after transfection, the growth media was exchanged for serum free DMEM and cells were incubated at 37°C and 5% CO_2_ for 1-2 hours. To image, the serum free DMEM was exchanged for room temperature phosphate-buffered saline containing 1 mM CaCl_2_ and 0.55 mM MgCl_2_. The plates were transferred to the microscope (Leica SP8X) and cells were imaged using a 40X oil objective with the following settings: 485 excitation and 525 emission wavelength, 5% laser power, HyD hybrid detector and a scan speed of 200 lines Hz (0.388 frames per second) with bidirectional scanning. All treatments were made in 1 mg/mL BSA in PBS and added directly to buffer for a final concentration of 0.1 mg/ml BSA.

### Gel-Based Activity-Based Protein Profiling

Membrane proteome was thawed on ice and 10 µg of protein was used for assay with volume normalized using 20 mM HEPES. Protein was incubated with activity-based probes, either 250 nM ActivX TAMRA-FP (ThermoFisher Scientific, 88318) or 500 nM MB064 (gift from Dr. Mario van der Stelt) for 15 minutes at 37°C. Reaction was quenched with 4X Laemmli Sample Buffer with 10% β-mercaptoethanol (BioRad) and run on a 10% polyacrylamide gel (Biorad). After running for about 1 hour at 150 mV, gel was removed from casing. Fluorescence was detected using a Chemidoc MP (Biorad) using a Cy3, green epifluorescence filter (605/50) for the activity-based probe and Cy5, red epifluorescence filter (695/55) for the protein ladder. The gel was then stained with Coomassie to obtain total protein to normalize results.

### 96-well Plate Reader GRAB_eCB2.0_ Detection

Clear-bottom, black 96-well plates (USA Scientific 5665-5087) were coated with poly-D-lysine (50 ng/ml, Sigma, P6407) for 1-2h at 37°C, after which the poly-D-lysine was removed, and coverslips were washed 3 times with sterile water and one time with DMEM. N2a cells were detached and resuspended in supplemented DMEM as described above, counted using a hemocytometer, then plated (20,000 cells per well) and were transfected after 24h with 0.1 µg DNA and 0.3 µg of PEI in 10 µl of serum free DMEM. 24h after transfection, growth media was initially replaced with serum free DMEM for 1h (with or without ABHD6 and DAGL inhibitors), before replacing with PBS supplemented with 1 mM CaCl_2_ and 0.55 mM MgCl_2_. Cells were incubated at room temperature for 20 min then a 1 min baseline fluorescence reading was obtained using a fluorescent plate reader using 485 emission and 525 emission filter settings with a 515 nm cutoff, and a speed of 1 reading every 20 sec. Immediately after baseline reading, treatments (made in 1 mg/mL BSA and PBS) were added to buffer in wells. Approximately 2 min after addition of treatment, the plate was read with same filter settings for 30 min. For cells pre-treated with SR1, SR1 made in PBS was added to cells after media had been replaced with PBS and were incubated for 20 min before baseline reading.

### LC-MS/MS

When N2A cells reached 90-100% confluency, the growth media was removed and replaced with serum free DMEM. Cells were incubated for 1 h at 37°C, after which the media was replaced with PBS supplemented with 1 mM CaCl_2_ and 0.55 mM MgCl_2_, incubated at room temperature for about 10 min and then treated for 2- or 10-min. Reaction was quenched by removing buffer and washing three times with ice cold PBS before scraping cells off and centrifuging at 500 x g for 10 min at 4° C to pellet the cells. The buffer was then aspirated, and the pellet resuspended in 20 µl of 0.02% TFA and 100 µl acetonitrile with 1 picomole of 2-AG-d^5^ internal standard on ice before transferring to 2.5ml of acetonitrile. Samples were briefly vortexed and incubated overnight at -20°C. After the overnight incubation, the homogenate was centrifuged at 2000 x g for 5 min to remove debris, the supernatant was collected and evaporated under nitrogen stream at 35°C and then resuspended in 50 µl of acetonitrile. The samples were capped under nitrogen stream and stored at -80°C until ready for the LC-MS/MS. Chromatographic separation was achieved using a Zorbax C18, 2.1 × 50 mm, 3.5 um reverse-phase column (Agilent). The HPLC output was directed into the electrospray ionization source of a Waters Xevo TQ-S mass spectrometer. Ionization was done in positive mode. All experiments were done in duplicate; results from each technical replicate were averaged and plotted using GraphPad Prism.

### Data Analysis

MATLAB, FIJI ImageJ, GraphPad Prism.

