## Supplementary Figures for "ABHD6 selectively controls metabotropic-dependent increases in 2-AG production"

### ABHD6 selectively controls stimuli-dependent increases in 2-AG levels

#### Supplementary Figures

**Figure S1:** Ponceau stain of Figure 1D

**Figure S2:** Concentration-dependent activation of $GRAB_{eCB2.0}$ by $CB_1R$ agonists.

**Figure S3a:** Measuring 2-AG, AEA, and AA by LC-MS: standards and N2a cell lysate.

**Figure S3b:** Ionomycin (2.5 µM) increases 2-AG levels in N2a cells measured by LC-MS and $GRAB_{eCB2.0}$ fluorescence.

**Figure S4:** Interpolation of 2-AG concentration, D034 activity and $GRAB_{eCB2.0}$ signals in Figure 4.

**Figure S5:** DAGLβ activity in N2a cells and KT-182 selectivity.

**Figure S1:** Ponceau stain corresponding to the detection of myc-$CB_1R$ or $GRAB_{eCB2.0}$ expression when transfected HEK293 and N2a cells using a western blot (see Figure 1D).

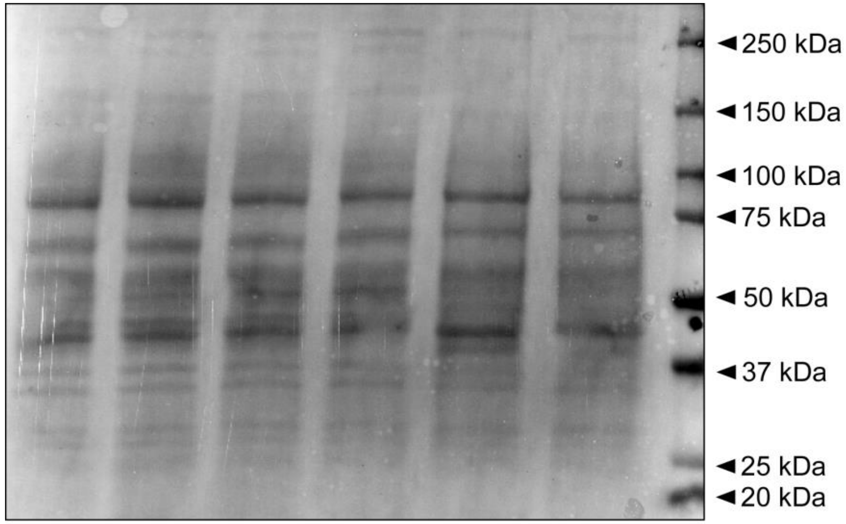

**Figure S2:** Concentration-dependent activation of $GRAB_{eCB2.0}$ by $CB_1R$ agonists. Direct activation of the $GRAB_{eCB2.0}$ in N2a cells was measuring using a fluorescent plate reader for 30 min (see Figure 2A). **A-B, D-G]** Time courses of fluorescent changes following treatment with increasing concentrations of CP (a), AEA (b), 1-AG (d), THC (e), Gly (f), and AA (g). **C]** Chemical structures of 1-AG and THC. **H]** Expression of mut-$GRAB_{eCB2.0}$ confirmed by immunostaining N2a cells transfected with mut-$GRAB_{eCB2.0}$ (i) or vector (ii) with a $CB_1R$ antibody. Time course data is shown as a mean of 3-10 independent experiments for AEA, 1-AG, THC, Gly, and AA and 3-39 independent experiments for CP. Experiments were done in triplicate; shaded area represent SEM. Gray area represent selected maximal response for mechanistic analyses. Scale bar in microscopy images represents 20 μm.

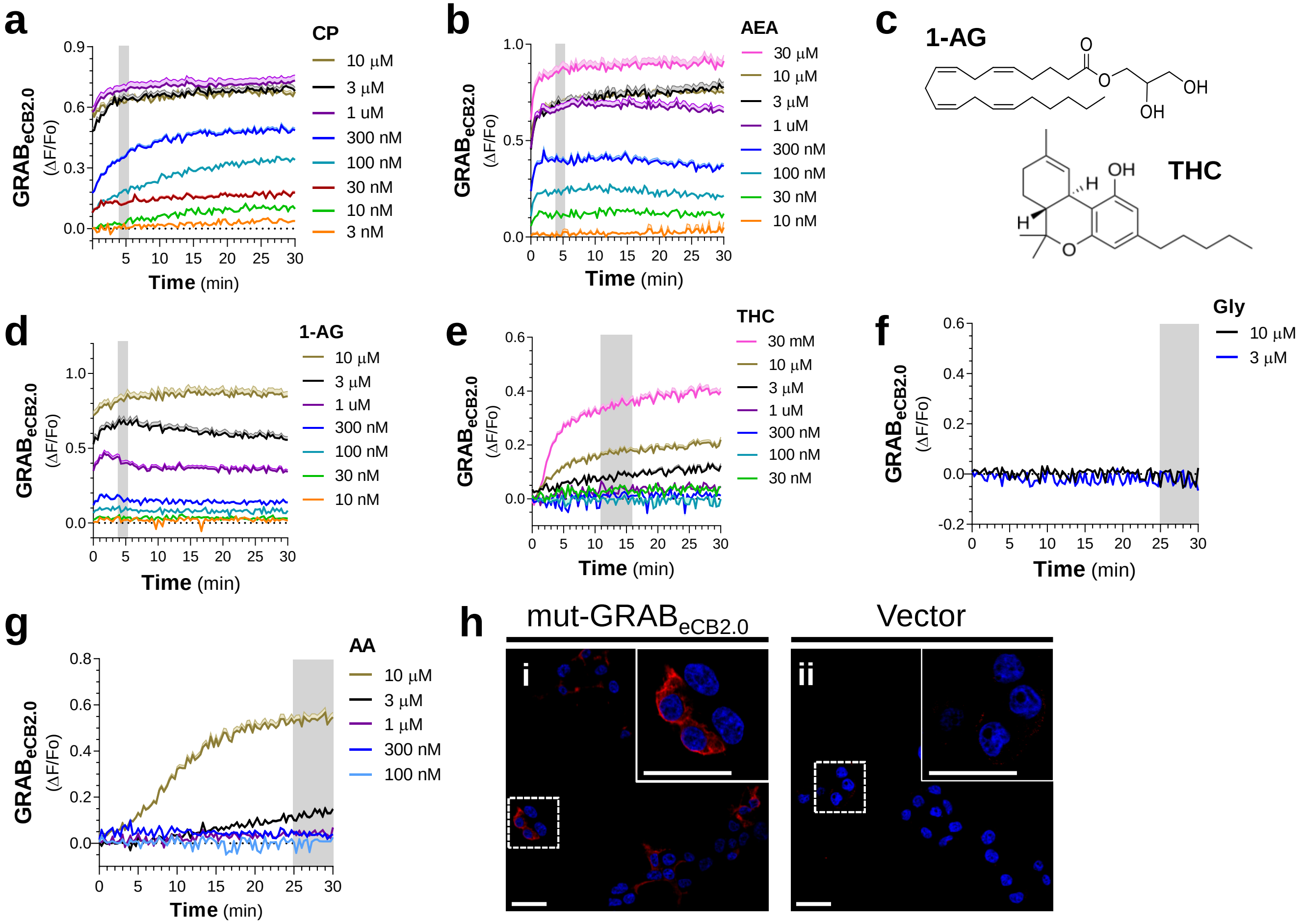

**Figure S3a:** Measuring 2-AG, AEA, and AA by LC-MS: standards and N2a cell lysate. **a, d, g]** LC-MS spectra of target analyte precursor ions obtained by injecting standards of 2-AG (a, *m/z* 379.35), AEA (d, *m/z* 348.43), and AA (g, *m/z* 303.23) standards containing internal standard of 2-AG-$d_5$ (*m/z* 384.35). **b, e, h]** Chromatograms of 2-AG (b), AEA (e), and AA (h) standards. Mass spectra and chromatograms of standard were generated by using 1 pmol of 2-AG, 1 pmol of AEA, and 5 pmol AA. **c, f, i]** Chromatograms detecting 2-AG, AA, and AEA in lysate made from vehicle treated N2a cells. Analytes were measured by monitoring the *m/z* transitions from 379.35 to 287.3 for 2-AG, 348.43 to 62.25 for AEA, and 303.23 to 205.13 for AA following collision-induced dissociation. All standard and N2a samples contained 0.2 pmol of 2-AG-$d_5$. The x-axis of chromatograms represents retention time in min. **j-l]** Representative images of the baseline fluorescence of a $GRAB_{eCB2.0}$-expressing N2a cell and the effect of mastoparan (10 μM) on cell fluorescence and morphology after 10 and 15 min of treatment (k and l respectively) as measured by live-cell confocal microscopy. Scale bar represents 10 μm.

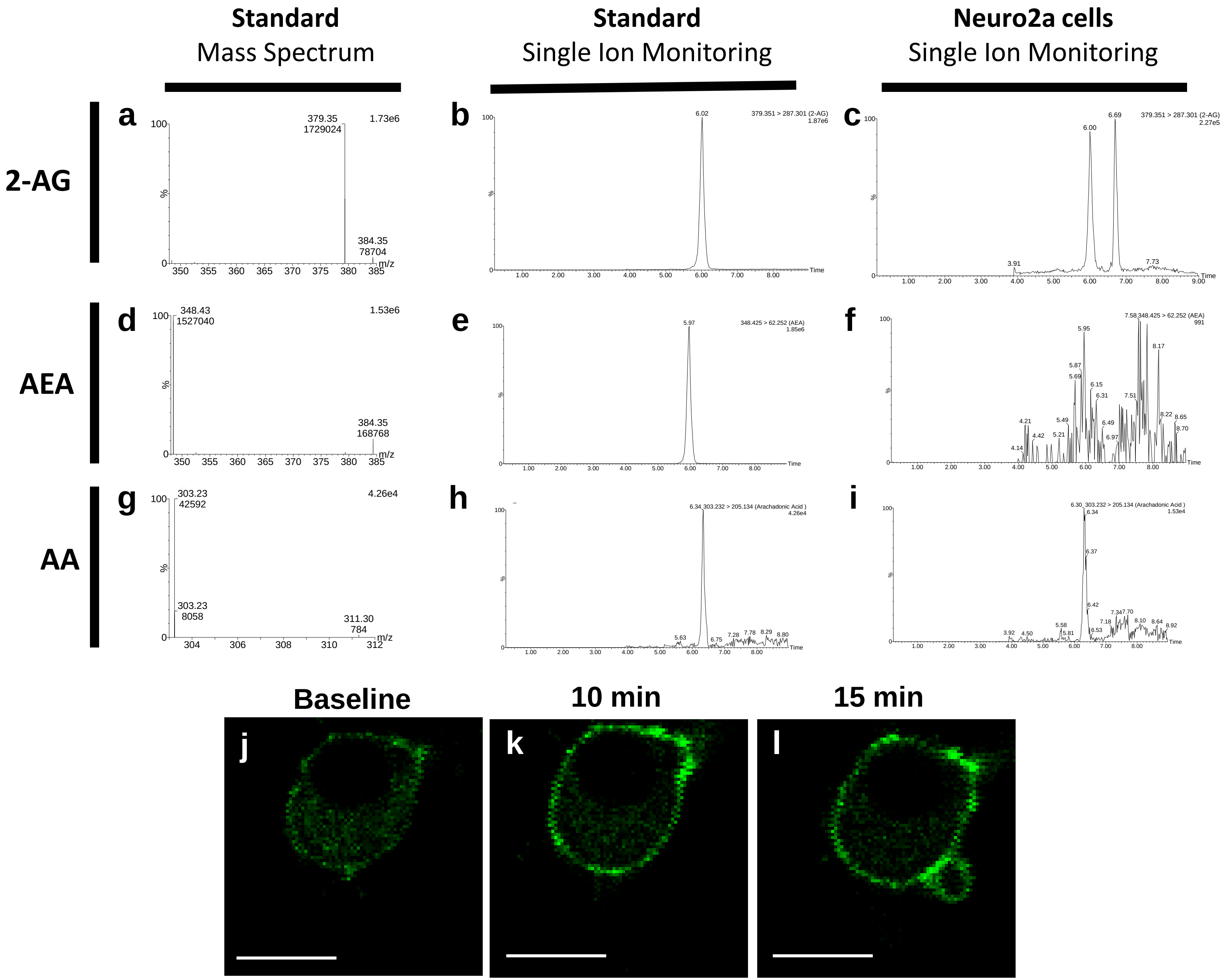

**Figure S3b:** Ionomycin (2.5 µM) increases 2-AG levels in N2a cells measured by LC-MS and $GRAB_{eCB2.0}$ fluorescence. **a]** Time-course of the effect of ionomycin on $GRAB_{eCB2.0}$ fluorescence in N2a cells as measured by live-cell confocal microscopy. Shaded area and error bars represent S.E.M. **b]** N2a cells were treated with ionomycin for 2 and 10 min and 2-AG levels measured by LC-MS/MS. Live-cell microscopy n = 35 cells from 3 independent experiments. LC-MS: vehicle n = 14; 2 min n = 10, 10 min n= 8. Significantly different from Vehicle (Veh): ANOVA followed by Dunnett's. *$p < 0.05$.

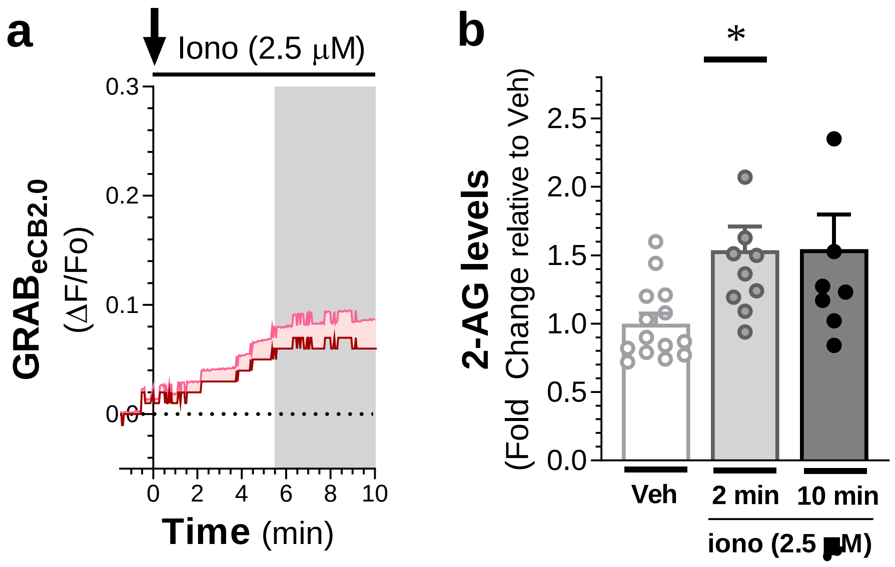

**Figure S4:** Interpolation of 2-AG concentration, D034 activity and $GRAB_{eCB2.0}$ signals in Figure 4. **a]** Interpolation of 2-AG concentrations following treatment of $GRAB_{eCB2.0}$-expressing N2a cells with BK (1 µM) or ATP (300 µM) (see Figure 4B, D) by using the concentration response curve generated by treatment with increasing concentrations of 2-AG (see Figure 2D) (log(agonist) vs. response -- Variable slope: Hill slope (0.45)). **b]** The DAGL inhibitor D034 has a concentration-dependent effect on DAGLβ inhibition in N2a cells. N2a cells were treated in situ with increasing concentrations of D034 (30 min, 37°C) followed by gel-based ABPP using the probe MB064 (1 µM, 15 min, 37°C). **c-d]** HOE, A74, and D034 do not affect $GRAB_{eCB2.0}$ fluorescence in N2a cells; however, removing extracellular calcium from the buffer (- Calcium) and TG decreases $GRAB_{eCB2.0}$ fluorescence. N2a cells expressing the mutant- $GRAB_{eCB2.0}$ experience a small and gradual increase in fluorescent signal following vehicle treatment. **e]** The direct activation of the $GRAB_{eCB2.0}$ by treatment with 2-AG (1 µM) is not affected by HOE, A74, D034, EGTA, TG, or by the removal of extracellular calcium. **f]** Mastoparan increases fluorescence in $GRAB_{eCB2.0}$ -expressing N2a cells in a concentration-dependent manner. In $GRAB_{eCB2.0}$ experiments, the fluorescent signal was measured using a fluorescent plate reader for 30 min, error bars represent SEM, N = 3-17 independent experiments done in triplicate.

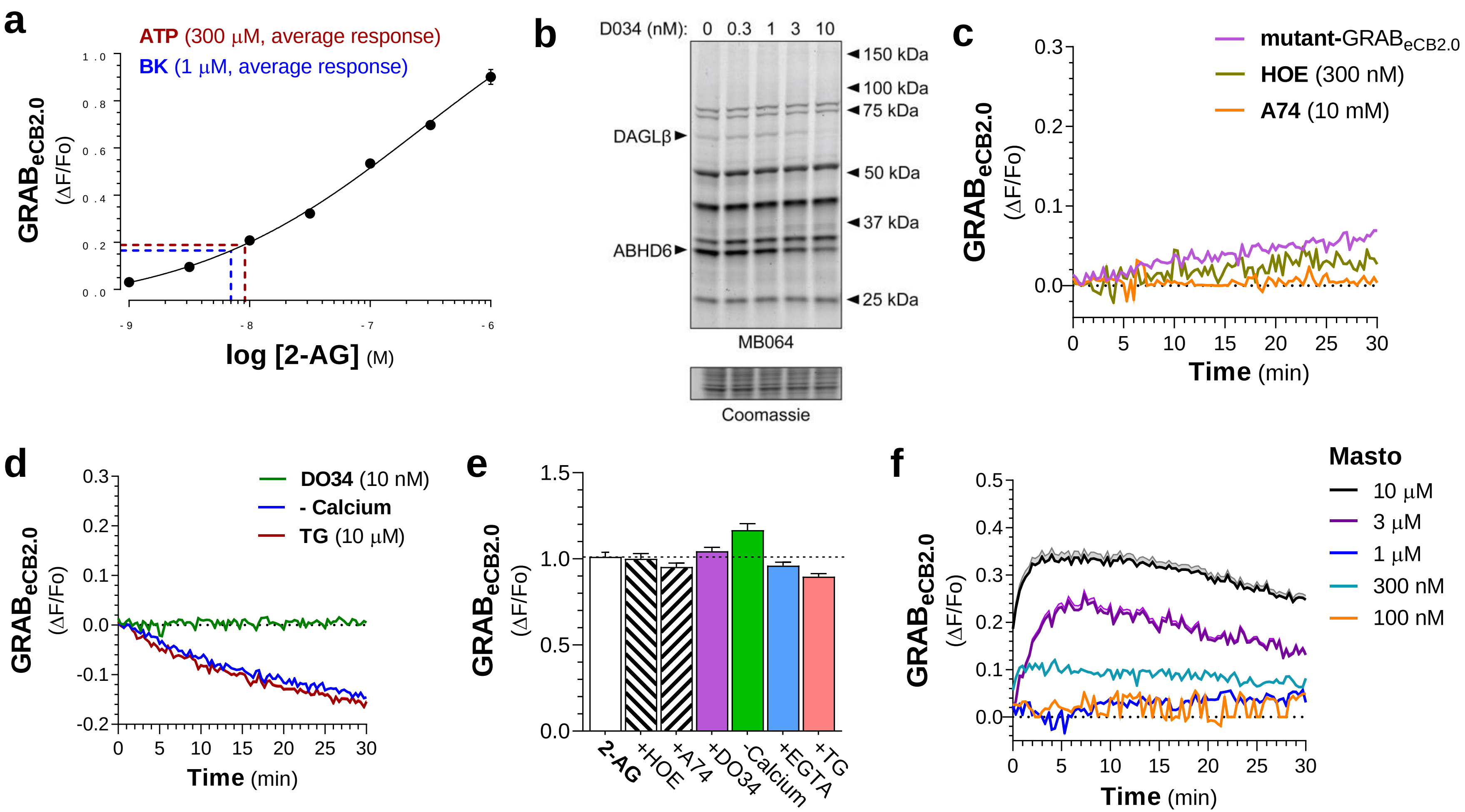

**Figure S5:** DAGLβ activity in N2a cells and KT-182 selectivity. **a]** N2a cells express DAGLβ activity, but not DAGLα activity, in contrast to mouse cortex (CTX) which express both DAGLα and DAGLβ activities. **b]** Representative image of gel showing that KT-182 inhibits ABHD6 in a concentration-dependent manner in N2a cells treated in situ for 60 min (see Figure 5B). **c]** KT-182 (10 nM, 60 min in situ) is selective for ABHD6 and does not significantly inhibit other enzymes labeled by the activity-based probes MB064 and TAMRA-FP (see Figure 5C). Cells were treated in situ with KT-182 and enzyme activity was measured in the membrane proteome of these cells by gel-based ABPP using either (c) MB064 (2.5 µM, 15 min, 37°C) or (d) TAMRA-FP (500 nM, 15 min, 37°C)

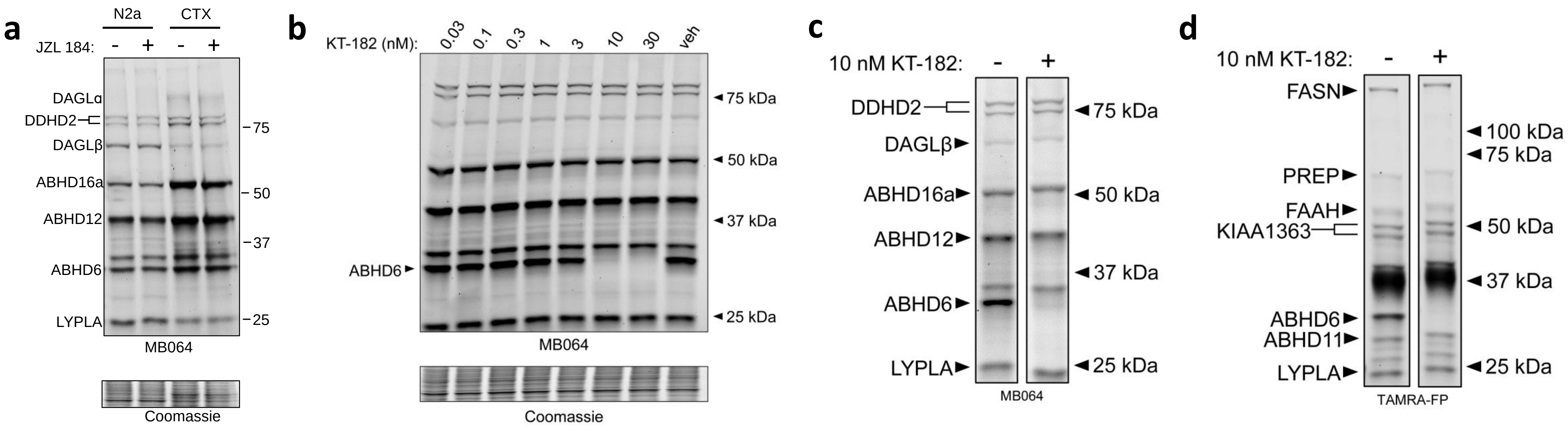
